# Nucleolar accumulation of APE1 through condensates is mediated by rRNA forming G-quadruplex structures

**DOI:** 10.1101/2024.03.04.583447

**Authors:** Giuseppe Dall’Agnese, Nancy M. Hannett, Kalon J. Overholt, Jesse M. Platt, Jonathan E. Henninger, Asier Marcos-Vidal, Giulia Antoniali, Gianluca Tell

## Abstract

APE1 (apurinic/apyrimidinic endodeoxyribonuclease 1) is the main endonuclease of the base excision repair (BER) pathway acting on abasic (AP)-sites in damaged DNA. APE1 is an abundant nuclear protein with a higher concentration than other BER pathway enzymes, and therefore, improper expression and localization of this factor could lead to the accumulation of toxic DNA intermediates. Altered APE1 sub-cellular localization, expression levels, or hyper-acetylation are associated with cancer development suggesting the importance of a fine-tuning mechanism for APE1 nuclear-associated processes. Recent work highlighted multi-functional roles of APE1, including rRNA quality control. However, how rRNA influences the sub-cellular localization and activity of APE1 remains poorly understood, but previously underappreciated APE1-RNA interactions may influence the ability of this protein to form biomolecular condensates and tune APE1 partitioning into nucleoli. Since nucleolar accumulation of ectopic proteins could be the result of overexpression strategies, it is imperative to have cellular models to study APE1 trafficking under physiological conditions. Here we created the first cell line to express fluorescently tagged APE1 at its endogenous locus, enabling live-cell imaging. Live-cell imaging demonstrates that APE1 nucleolar accumulation requires active rRNA transcription. When modeled in vitro, APE1 condensate formation depends on RNA G-quadruplex (rG4) structures in rRNA and is modulated by critical lysine residues of APE1. This study sheds light on the mechanisms underlying APE1 trafficking to the nucleolus and formation of RNA-dependent APE1 nucleolar condensates that may modulate a switch between the activity of this factor in rRNA processing and DNA damage repair.

**Significance Statement:** We created and characterized the first endogenous, fluorescently tagged cell line to study APE1 subcellular trafficking under physiological and stress conditions. Using this cell line, we show that APE1 nucleolar enrichment occurs under physiological conditions and, performing *in vitro* droplet assays, we associate APE1 condensates with active transcription of RNA G-quadruplexes, abundantly present in healthy nucleoli. This work deepens our understanding of APE1’s role in healthy cells in the absence of DNA damage and provide a novel mechanism for how this protein responds to stress. Our results suggest that phase separation is an important part of how DNA damage repair proteins switch between their normal physiological functions and their ability to correct DNA lesions.

## Introduction

Damage to DNA is a threat to the cell and different DNA damage response (DDR) pathways have been acquired during evolution to cope with insults to DNA, which involve the coordinated action of different proteins and enzymes as well as the generation of several toxic intermediates that should be promptly and efficiently repaired in order to preserve genomic stability and to avoid the activation of the death signaling cascades (1). APE1 (apurinic/apyrimidinic (AP) endodeoxyribonuclease 1) is an enzyme discovered for its central role as the main endonuclease acting on the base excision repair (BER) pathway on abasic (AP)-sites generated spontaneously or by the action of several glycosylases (2, 3). APE1 is also known for its other, non-canonical, functions such as redox-regulated activity on different transcription factors (4), miRNA processing (5, 6) and RNA quality control (7, 8). APE1 is an abundant protein within human cells ranging a concentration from about 0.2 to 10 μM (9), which is up to 100-fold higher than the downstream enzymes in the BER pathway (10) and, therefore, improper expression of the protein could generate the accumulation of toxic DNA intermediates, mainly single strand breaks (SSBs). The APE1 C-terminus, responsible for the endonuclease activity through which it generates SSBs upon recognition of the abasic sites, is highly conserved during evolution, meanwhile its N-terminus (comprising residues 1-33), responsible for the non-canonical functions of the protein, seems to be recently acquired during evolution (11). In cancer cell lines, APE1 shows nuclear localization with enrichment in the nucleolus due to NPM1 nucleolar localization under overexpression conditions (7, 12). Several pathologies, mostly cancers, are associated with either altered APE1 sub-cellular localization (13–15), expression levels (mostly over-expression) (16–18), or hyper-acetylation (19, 20), suggesting the importance of a fine-tuning mechanism for APE1 nuclear-associated processes.

Our lab has previously shown that APE1 endonuclease activity participates in rRNA quality control and nucleolar enrichment through the interaction with NPM1 (15, 21–25). Biochemical purification of nucleoli showed that APE1 nucleolar enrichment, along with other BER-associated proteins, is highly regulated by NPM1; either NPM1 knockdown or its subcellular re-localization upon cisplatin treatment leads to APE1 nucleolar depletion (24). We previously characterized several APE1 acetylated mutants by using cellular reconstitution assays: HeLa cell clones able to express a specific, inducible, APE1-siRNA to knock down the endogenous protein and re-express ectopic FLAG-tagged forms through siRNA-resistant cDNA expressing plasmids. With this cellular model, we discovered that acetylation of lysine K^27^/K^31^/K^32^/K^35^ occurs upon genotoxic stimuli and, in addition to being relevant for APE1-NPM1 interaction, plays essential roles in both APE1 DDR-associated enzymatic activity (21) and nucleic acid interactions (10, 23). Motivated by our previous works, several labs have investigated the role of acetylation on lysine at the APE1 N-terminal region, demonstrating the interconnection between various regulatory factors (26). Additionally, acetylation of APE1 N-terminal lysines seems to play a newly discovered role in the binding and stabilization of DNA G-quadruplex (G4) complexes. APE1 can bind G4s through its N-terminal domain, and acetylation of lysines stabilizes those complexes by increasing APE1 residence time (27–30). Until now, a major limitation of all the studies concerning APE1 is that they have been performed using overexpression cell models, therefore there is an urgent need for additional cell models to characterize the functions of the protein under more physiological conditions. This work addresses this issue by providing definitive support to our previous findings and proposing a new mechanism for the observed nucleolar enrichment of the protein due to RNA G4-mediated condensates.

Compartmentalization of specific biochemical reactions in DDR condensates is emerging as a novel mechanism to efficiently coordinate DDR (1). Condensates, defined as dynamic organelles not enclosed within membranes, are subcellular compartments mostly formed by protein and nucleic acids (31). Among the different ways through which cells can regulate condensates, post-translational modifications (PTMs) are known to play important roles since they can either promote or suppress a protein’s phase separation (e.g. acetylation is known to disrupt condensates) (32, 33). The rising importance of understanding the physicochemical properties of condensate formation and the ability to compartmentalize potentially harmful biological reactions (34) could be important for developing new therapeutic approaches (31, 35). For instance, condensation mechanisms have been recently described in the case of: i) the RNA-dependent recruitment of 53BP1 and MRNIP-mediated coordination of MRN complex for DSB repair, leading to the resolution of the damage (36, 37); ii) the regulation of several DDR proteins (i.e. ERCC1 and EXO1) through SENP6-and SLX4-controlled SUMOylation, allowing DDR-associated proteins to avoid excessive protein turnover and degradation (38, 39) and iii) the APE1-driven phase separation promoting ATR-Chk1 damage response, in which APE1 nucleolar enrichment promotes recruitment of both ATR and its activators (TopBP1 and ETAA1), leading to ATRzmediated Chk1 phosphorylation (40). Common aspects of all those DDR proteins able to form or to be compartmentalized into DDR condensates include the presence of an intrinsically disordered region (IDR) and the ability to bind to RNA, two characteristics that help proteins undergo phase separation (41–45).

The nucleolus, one of the first condensates to be described (46), plays an important role in ribosome biogenesis, particularly in ribosomal RNA (rRNA) synthesis and processing (47). Nucleoli possess a very peculiar organization, divided into three mutually immiscible, liquid phases (48). Recent evidence suggests that ribosomal DNA (rDNA) could form G-quadruplexes (G4) (49), structures also observed in rRNA, forming RNA-G4 (rG4) (50). rG4s are known to play different roles in cellular biology, ranging from telomere protection, regulation of transcription, inhibition of translation, and replication-dependent genome instability (51). Interestingly, rG4s is that they seem to play a role also in condensate formation (52, 53). Considering that rRNA is predicted to fold into several rG4 structures, it might be possible that the other nucleolar functions can occur through rG4s; nucleoli play a fundamental role as: i) a “stress sensor” by regulating the p53 signaling pathway (54); ii) a source of transcription of different RNA molecules beside rRNA (55), and iii) a hub for maintenance of genome stability since they are able to coordinate and accumulate over 150 different DDR-associated proteins (56). All those non-canonical nucleolar-associated mechanisms have been summarized in (57).

Recently, Yan and colleagues, using overexpression cancer cell models, showed that APE1 can undergo phase separation promoting ATR-Chk1 condensate formation in the nucleolus, emphasizing that APE1’s IDR is essential for condensate formation (40). ATR plays important roles in regulating cell cycle checkpoints as well as the homologous repair (HR) pathway and a recent study demonstrated that APE1 overexpression is involved in the dysregulation of the HR pathway (18), suggesting that a better model, with physiological expression of APE1, is needed to investigate APE1’s ability to undergo phase separation in living cells under normal conditions.

In this work, we generated and deeply characterized what, to our knowledge, is the first cell line with a fluorescently tagged APE1 endogenous gene. Using this model, we were able to follow the dynamics of APE1 subcellular trafficking under basal conditions and different genotoxic treatments, showing that nucleolar enrichment of the protein was dependent on active rRNA transcription and could be modulated by cisplatin (CDDP) treatment. Moreover, using purified recombinant proteins, we assessed the influence of DNA and RNA on APE1 protein droplets *in vitro*, showing that rG4s present in rRNA genes stimulate APE1 condensate formation.

## Results

### Generation and characterization of endogenous APE1 mEGFP-tagged mESCs

To visualize APE1 dynamic movements within living cells, avoiding non-specific effects due to overexpression strategies, we tagged APE1 at the endogenous locus with mEGFP at either its N- or C-terminus, using the CRISPR/Cas9 system in mESCs (schematic representation of the model used in Fig.1A). To confirm successful tagging and to select cells tagged in a homozygous manner, we picked 32 colonies and confirmed successful tagging via PCR on isolated genomic DNA (Supp. Fig.1A). From these, 5 homozygously tagged clones were selected for downstream analyses (Supp. Fig.1A). To observe if the tagging had any effect on APE1 subcellular localization, we performed live cell-imaging analyses on the selected clones. Interestingly, we observed that N-terminus tagged APE1 showed an altered nucleolar enrichment and, therefore, decided to proceed only with the validation and characterization of the C-terminus tagged clones (Fig.1B and Supp. Fig.1B).

**Figure 1.**
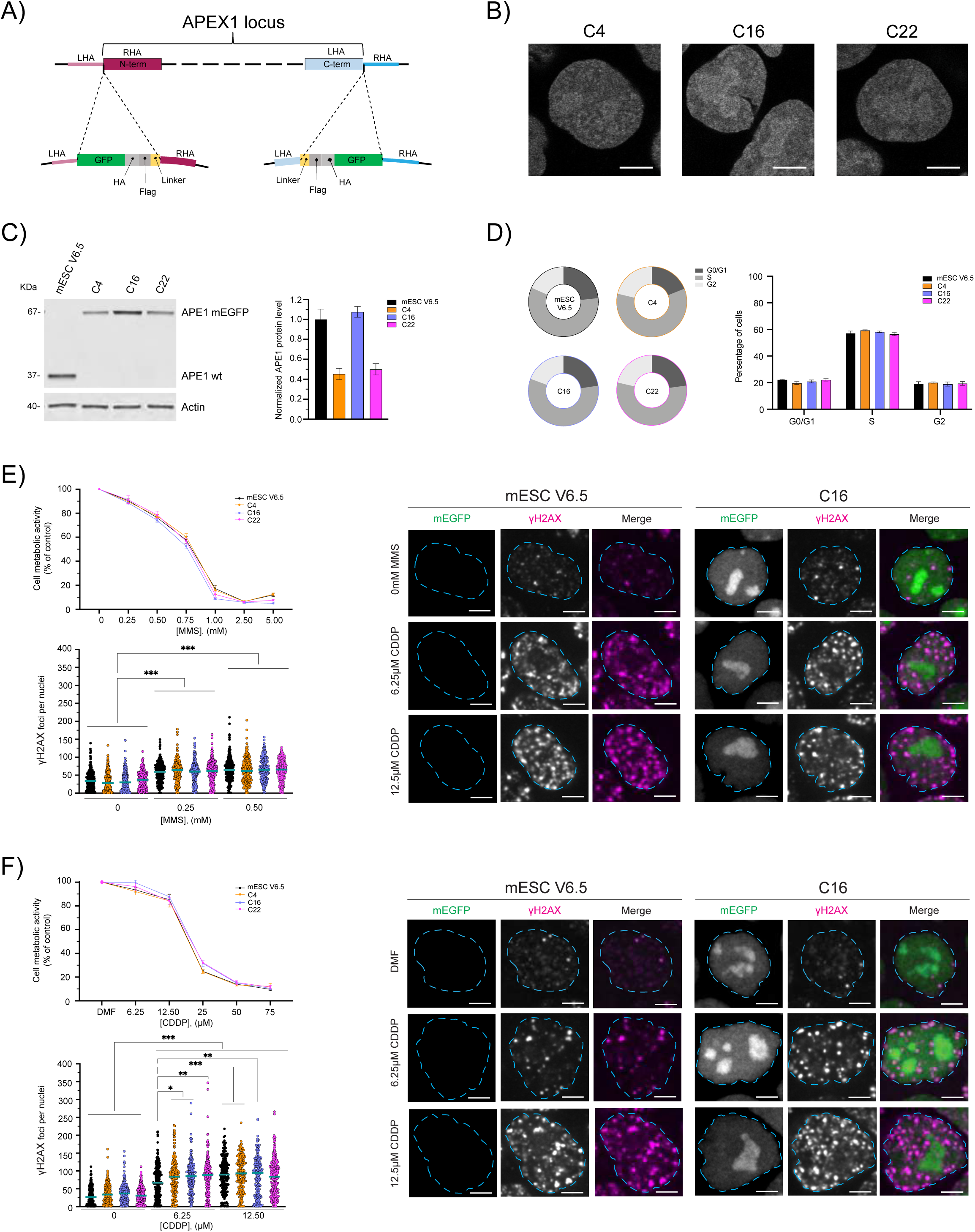
Generation and characterization of APE1 endogenous tagged murine Embryonic Stem Cells (mESC). Schematic representation of CRISPR-Cas9 technology used to generate the endogenously tagged mESC; N-terminus tagging of APE1 to the left and C-terminus tagging to the right (A). Live cell imaging of C4, C16, and C22 clones with APE tagged at the C-terminus of the protein show proper nuclear localization, with nucleolar enrichment of APE1; images taken with a 63x objective of a Zeiss LSM 980 with Airyscan 2 Laser Scanning Confocal microscope, scale bar 5 μm (B). Representative western blot, with respective quantification analysis, quantification performed on a biological triplicate and normalized by Actin protein level setting APE1 wt as 1; error bars represent SEM (C). Cell sorting for cell cycle analysis on APE1 tagged clones, pie charts representing a single experiment are shown to the left, and a histogram with biological triplicates is shown to the right; error bars represent SEM (D). MTS viabilities assays of cells treated with different doses of MMS for 8 hours (E) or CDDP for 24 hours (F), values of four biological replicates have been normalized based on control (plain media or media containing DMF respectively); error bars represent SEM. Staining for γH2AX on cells treated either with control (plain media), 0.25 mM or 0.5 mM MMS for 8 hours (E) and control (DMF), 6.25 μM and 12.5 μM CDDP for 24 hours (F) with γH2AX foci count and representative images of wt and C16 clone are shown; images taken with 100x objective of an RPI Spinning Disk Confocal microscope, foci measurement performed on biological triplicate (2 way ANOVA statistical analysis with * - p<0.05, ** - p<0.001, *** - p<0.0001).

We then evaluated whether the addition of the tags might alter protein expression levels. By performing Western blotting analyses, with a specific anti-APE1 antibody, we observed that one out of the three selected clones (C16) possessed comparable protein levels to the wild-type cells, whereas the other two clones (C4 and C22) showed almost half of the protein amount with respect to the wild-type cells (Fig.1C). As an orthogonal approach, we assayed protein expression level using FACS analysis, which demonstrated that clones C4 and C22 clones had half the expression of APE1 when compared to clone C16, consistent with the results of the western blot analysis (Supp. Fig.2B).

As a following step, we assessed if the expression of the engineered tagged-APE1, or the lower expression levels, might alter the cell cycle distribution of mESCs. We performed a cell cycle analysis on sorted cells and concluded that neither the addition of the tag nor the decrease in APE1 amounts had a major effect on the G0/G1, S, and G2 distribution of the cell cycle (Fig.1D and Supp. Fig.2A).

To confirm that endogenous tagging did not alter APE1 function, we assayed the function of APE1 in wildtype and tagged cells. APE1 is mostly known for its DNA damage repair ability, especially in repairing abasic sites, generated spontaneously or as a consequence of glycosylase action on oxidized alkylated bases, via the BER pathway (3, 9, 58). For this reason we treated cells with different doses of methyl methanesulfonate (MMS), a well-known alkylating agent, which induces DNA lesions specifically repaired by the BER pathway (11, 59, 60). We also used CDDP, which generates lesions that are also repaired by APE1, though independent of the BER pathway (17, 61–63) (Fig.1E and 1F). We first evaluated cell viability via an MTS assay of wild-type cells and the selected clones, which showed no significant alteration in cell toxicity (Fig 1E and 1F). To further evaluate the efficiency of the DNA damage response of the clones, we measured the number of γH2AX foci generated upon low doses of MMS (0.25 and 0.5 mM) or CDDP (6.25 and 12.5 μM) and noticed similar treatment responses comparable to the wild-type cells (Fig.1E and 1F).

Altogether, these results show that the addition of the tags at the N-terminus altered APE1 subcellular localization, whereas the C-terminal tagging did not alter either APE1 subcellular localization or DDR activity. More importantly, we developed the first non-tumoral cell model based on the expression of the endogenous APE1 protein fused with mEGFP, which allows monitoring of subcellular trafficking under physiological, non-overexpressing conditions.

### APE1 nucleolar accumulation requires active rRNA transcription as determined from live cell imaging analyses

We tested whether treatment with the DNA-damaging compounds MMS and CDDP may affect APE1 subcellular distribution. By treating cells with 0.5 mM of MMS, we observed no alteration of APE1 subcellular localization within the 6 hours of acquisition (Fig.2A, Supp. Fig.3A and Supp. Fig.3B). On the contrary, treating cells with 50 μM of CDDP for 6 hours was able to induce APE1 nucleolar depletion even at short times, with statistically significant differences starting at 180 minutes of treatment (Fig.2B, Supp. Fig.3C and Supp. Fig.3D). To test whether the effect of CDDP is more likely to be related to its DNA platinating activity or its ability to inhibit RNA Pol I transcriptional activity (64–67), we treated cells with 1 μM of CX5461, a clinically used RNA Pol I inhibitor known to bind to and stabilize G4 structures (68). As expected, we observed the same APE1 re-distribution as in CDDP-treated cells, with the first statistically significant difference observable after 9 minutes from the start of the treatment (Fig.2C, Supp. Fig.3E and Supp. Fig.3F). These results show that MMS-induced DNA damage alone is not sufficient to drive APE1 exit from the nucleolus, whereas inhibition of rRNA transcription by RNA Pol I can cause APE1 to re-localize outside of the nucleolus.

**Figure 2.**
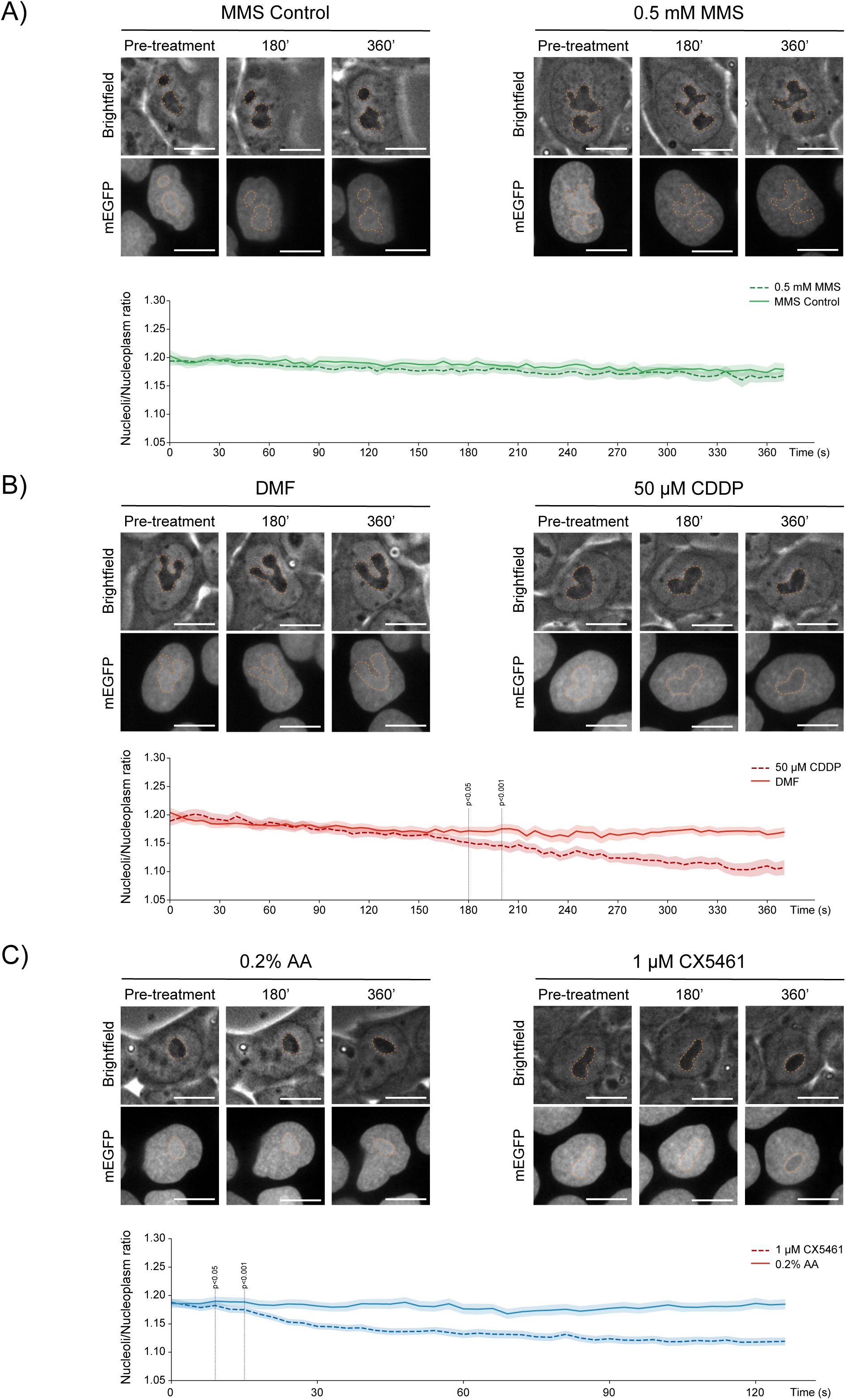
APE1 nucleolar localization is dependent on rRNA transcription. Live cell imaging of endogenously tagged APE1 with mEGFP taken with a 100x objective of Andor Revolution Spinning Disk Confocal, FRAPPA, and TIRF system microscope. Representative images of cells before (pre-treatment), at 180 and 360 minutes of treatment with either 0.5 mM MMS (A) or 50 μM of CDDP (B), with respective controls, are shown; nucleoli/nucleoplasm ratio of tagged APE1 signal over time are shown on the graph below the representative images; measurements acquired for at least 3 different fields of the same well with the experiment repeated 4 times, over 150 nuclei were considered for the analysis of each treatment. Representative images of cells before (pre-treatment), at 9 and 120 minutes of treatment with 1 μM of CX5461 (C) and its control (0.2% acetic acid) are shown; nucleoli/nucleoplasm ratio of tagged APE1 signal over time are shown on the graph below representative images; measurements acquired for at least 4 different fields of the same well with the experiment repeated 4 times, over 150 nuclei were considered for the analysis. Nucleoli are highlighted in orange dotted lines and scale bar of 5 μm.

### rG4 present in rRNA promotes APE1 droplet-forming ability, which is mediated by critical lysine residues

Considering the recently published evidence that APE1 forms condensates in the nucleolus, promoting DDR and that it forms droplets in an *in vitro* droplet assay (40), we investigated whether we were able to recapitulate those findings using a buffer normally used for APE1 enzymatic activity *in vitro* (AP-Buffer) (7, 62). Further, we assessed if different APE1 mutants (APE1^NΔ33^, APE1^K4pleA^, and APE1^K4pleR^) retain the ability to form *in vitro* droplets similar to the wild-type protein (APE1^WT^) (Fig.3). Using metapredict, a deep-learning consensus predictor of intrinsic disorder and protein structure (69, 70), we observed that APE1^WT^ possesses a 60 amino acid long IDR at its N-terminus ((1) and Supp. Fig.4A). Interestingly, deletion of the first 33 amino acids (APE1^NΔ33^) does not alter the APE1 disorder/order prediction in the remaining sequence (Supp. Fig.4B). We choose to investigate APE1^NΔ33^ since the deletion of this portion can be considered a naturally occurring PTM within the cytoplasm (71), and particularly, because the APE1 N-terminus is known to be essential for protein-protein interactions (7, 21, 26, 40, 72) (e.g. NPM1 and ATR) as well as RNA-binding (7, 10, 21). Among different PTMs, acetylation is known to impair a protein’s ability to undergo phase separation (73). For this reason, we decided to investigate whether APE1 mutants with neutralized positive charges, mimicking lysine acetylation, might impair its ability to form *in vitro* droplets using two mutants that have no alteration in the predicted IDR: APE1^K4pleA^ (Supp. Fig.4C) and APE1^K4pleR^ (Supp. Fig.4D). APE1^K4pleA^ and APE1^K4pleR^ are mutated in lysine positions 27, 31, 32 and 35 to alanine, mimicking the acetylated form of APE1, or to arginine, mimicking a non-acetylated form of APE1, respectively.

**Figure 3.**
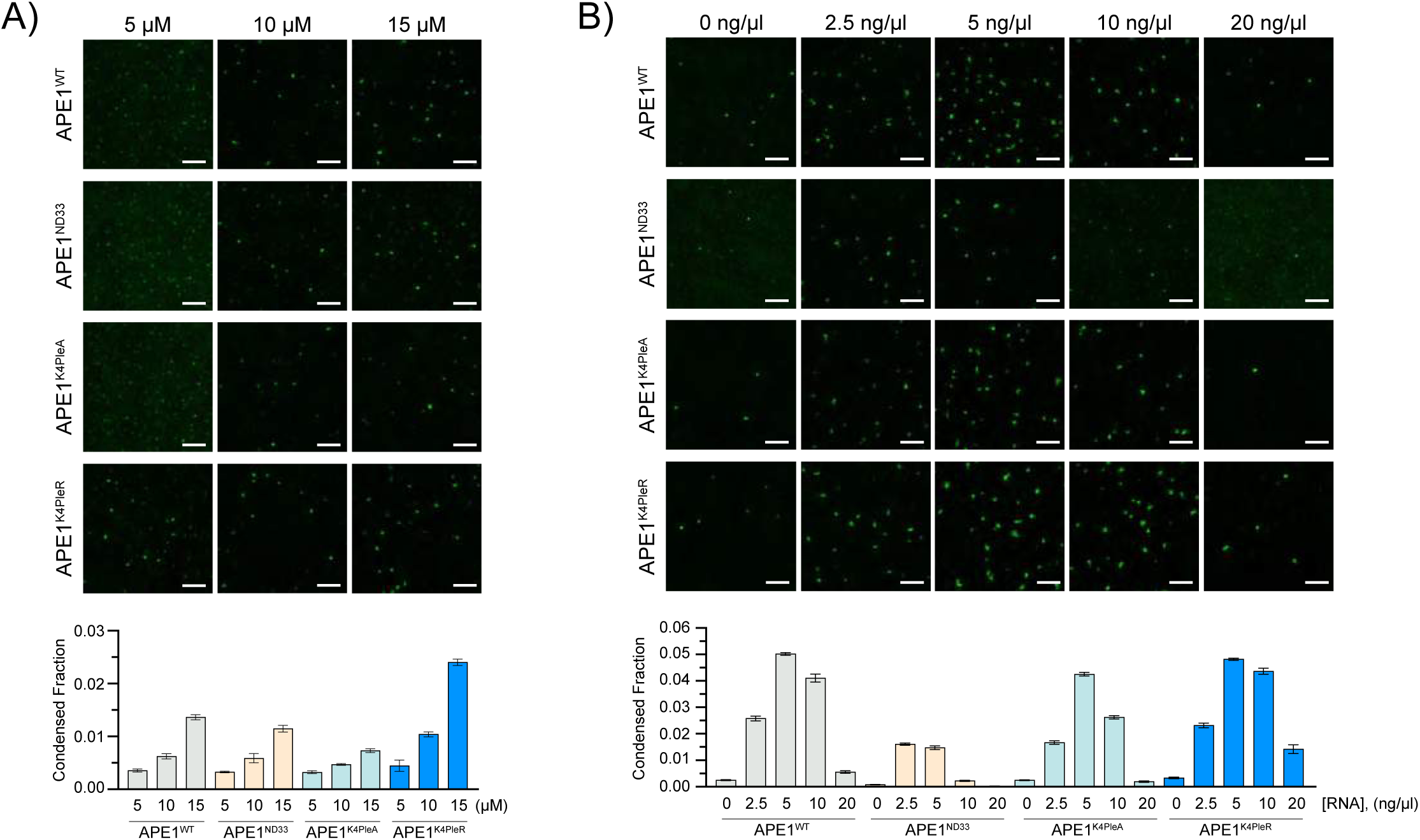
RNA influence on *in vitro* purified APE1-mEGFP protein droplet formation. Representative images (top) with background subtraction of in vitro droplet assay, with respective quantification (bottom) of APE1 droplets. A) 5, 10, and 15 μM of APE1 proteins (APE1^WT^, APE1^NΔ33^, APE1^K4pleA^, and APE1^K4pleR^) with respective measurements of condensed fraction. B) 10 μM of APE1 proteins (APE1^WT^, APE1^NΔ33^, APE1^K4pleA^, and APE1^K4pleR^) in combination with 0, 2.5, 5, 10, and 20 ng/μl of total RNA with respective measurement of condensed fraction. Every image shows a 5 μm scale bar. The experiment was performed twice with similar trends; images and analysis shown are of a single experiment. The experiment was performed in duplicate and showed the same trend; the images and analysis shown are of a single experiment.

By performing an *in vitro* droplet assay with different purified protein concentrations, we observed that all the proteins were able to form droplets within a range from 5 to 15 μM with APE1^K4pleR^ being the protein with the highest tendency to form droplets, followed by APE1^WT^. As expected, APE1^K4pleA^ and APE1^NΔ33^ demonstrated a reduced ability to form droplets *in vitro* when compared to APE1^WT^, especially at higher concentrations (Fig.3A). Besides protein concentration, another parameter that can influence the ability of a protein to form droplets *in vitro* is the presence or absence of RNA molecules (41, 43, 44, 74). For this reason, we extracted total RNA, which is mostly constituted by rRNA, from mESCs and performed *in vitro* droplet assays with increasing concentrations of RNA (Fig.3B). We observed that the ability to form droplets of all the proteins used was affected in a dose-dependent manner by the presence of RNA, with a maximum increase at 5 ng/μl of RNA. APE1^WT^ and APE1^K4pleR^, known to directly bind RNA (21), promoted better droplet formation when compared to APE1^K4pleA^ and APE1^NΔ33^. Surprisingly, the two mutants with impaired ability to bind RNA (i.e. APE1^K4pleA^ and APE1^NΔ33^) were also influenced, showing a higher condensed fraction in the presence of RNA (Fig.3B).

APE1 is mostly known as a DDR enzyme (3, 11, 58), therefore we wanted to assess whether DNA can partition into APE1 droplets. To do so, we used two DNA sequences, one containing the AP site-mimicking tetrahydrofuran (THF) and one unmodified control (respectively listed as THF and DNA in Supp. Table 2). Besides its DNA repair activity, APE1 is emerging as an important RNA processing protein (2, 5, 8, 75), particularly implicated in rRNA processing (7, 21). Considering our hypothesis that rRNA active transcription is the key element necessary to keep APE1 enriched within the nucleolus and that rRNA is predicted to form several rG4s (Supp. Fig.5), we tested the influence of a putative rG4 on droplet formation for all the APE1 proteins mentioned above. First, we assessed how many rG4s are predicted to form in the murine 45S rRNA; using QGRS Mapper, we found 120 sequences, 31 in the 5’ ETS, 8 in the 18S, 10 in the ITS1, 0 in the 5.8S, 13 in the ITS2, 49 in the 28S and 9 in the 3’ ETS (Supp. Fig.5 left table). We chose the highest 20-mer rG4 sequence from the 18S to investigate further (Supp. Fig.5 right table). We designed a Cy5-labelled probe with the predicted rG4 sequence and, as control, we used a 20-nucleotide long rPoly-U sequence (sequences shown in Supp. Table 2).

At first, to test whether those probes (dsDNA, dsTHF, rG4, and rPoly-U) might alter APE1’s ability to form droplets *in vitro*, and to select the best concentration to test with the mutants, we performed *in vitro* droplet assay, with increasing concentrations of those probes using APE1^WT^ (Fig.4A). Interestingly, we observed that dsDNA and dsTHF had only a mild influence on APE1’s ability to form *in vitro* droplets using concentrations ranging from 0.25 to 2 μM (Fig.4A top panel). The rPoly-U probe showed a similar influence on APE1^WT^, whereas the same concentration of rG4 RNA probes improved APE1’s ability to form *in vitro* droplets, even at concentrations as low as 0.25 μM (Fig.4A bottom panel). By monitoring the probe localization using the Cy5 label, we observed an enriched signal for each probe inside of APE1^WT^ droplets, suggesting that they partition into APE1 droplets. In support of our hypothesis, at similar probe concentrations, rG4 had a much higher ability to partition into APE1 droplets compared to all the other probes (Fig.4 quantification panels). Selecting 0.5 μM as the “optimal” probe concentration, we compared differences in droplet forming ability and partitioning of APE1^WT^ with the mutants mentioned above. Comparing APE1^WT^ with the mutants, we observed that: i) the presence of general RNA (rPoly-U in our case) particularly affects APE1^K4pleR^ droplet ability; ii) the dsDNA and dsTHF probes can partition similarly between the different mutants and iii) rG4 can partition in droplets formed by all the tested proteins, with a higher partition ratio in APE1^WT^ and APE1^K4pleR^ (Fig.4B). Altogether, these findings confirmed the ability of APE1 to form *in vitro* droplets in the AP-Buffer, and demonstrated improved droplet formation when proteins were co-incubated with an rRNA sequence predicted to form rG4.

**Figure 4.**
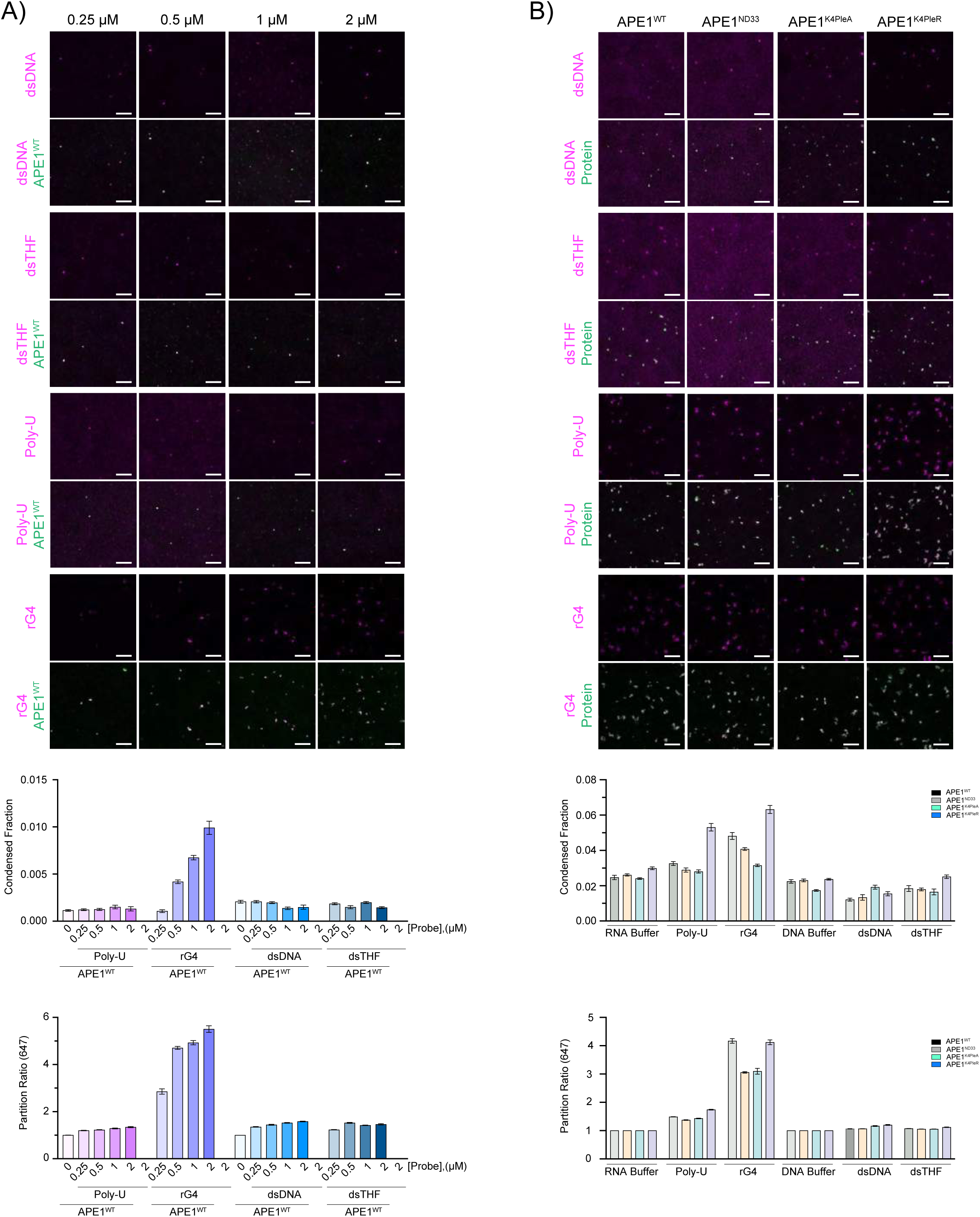
rG4 promotes *in vitro* purified APE1-mEGFP protein droplet formation. Representative images (top) with background subtraction of in vitro droplet assay, respective quantification of APE1 droplets and nuclei acid partition into APE1 droplets are shown in the bottom panels. A) 10 μM of APE1^WT^ with different concentrations (0, 0.25, 0.5, 1, and 2 μM) of probes (dsDNA, dsTHF, rPoly-U, and rG4), respective measurements of condensed fraction and partition ratio are shown. B) 10 μM of the different proteins (APE1^WT^, APE1^NΔ33^, APE1^K4pleA^, and APE1^K4pleR^) with 0.5 μM of the different probes (dsDNA, dsTHF, rPoly-U, and rG4), respective measurements of condensed fraction and partition ratio are plotted on the histograms. Scale bar of 5 μm. Images and analysis shown are of a single experiment. The experiment was performed in duplicate and showed the same trend; the images and analysis shown are of a single experiment.

## Discussion

The BER-associated endonuclease APE1 is essential for several biological functions, including DNA repair (3), RNA processing, and RNA quality control (76). APE1, the only protein within the BER pathway possessing endonuclease activity, is essential for proper cell homeostasis and its reduction or absence can drastically impair organismal viability and development (77–79). Until now, functional studies concerning the characterization of APE1 different functions and interactions have been only performed using overexpressing or knockdown/knockout cell models and mainly using cancer cell lines. Fine-tuning mechanisms are necessary to modulate the different functions of APE1 and it is emerging that their alterations are involved in the onset of pathological states. In particular, there is a growing amount of evidence that associates APE1 overexpression with the development of several types of cancers including lung (17), colon (80), liver (81), prostate (82), ovarian (83), and bladder cancer (84). In addition to that evidence, a recent study suggested that elevated APE1 protein levels are associated with increased genome instability, leading to cellular oncogenic transformation (18). The only cellular model that approaches physiological fidelity, although imperfectly, is an inducible model previously published by our lab (7). This model allowed us to regulate the suppression of endogenous APE1 while concurrently expressing the recombinant protein to a comparable degree (7). Along with the need for proper APE1 expression models to help us understand its physiological cellular functions, when considering the activity of proteins in the context of biomolecular condensation mechanisms, one must consider that protein concentration is a critical parameter (35, 58, 59). Due to these considerations, there is an urgent need to develop effective tools to study APE1 cellular activity under conditions closely mirroring cellular physiology. For these reasons, we decided to endogenously tag the APE1 gene in mESCs. The use of this cell model offers the additional advantage of its potential in studying APE1 functions under different conditions, throughout cellular differentiation, and, potentially, also at the organismal level due to the ability of mESCs to generate fully developed mice (85).

We observed that APE1, tagged at its C-terminus, retained its well-established nuclear localization with nucleolar enrichment (7, 12); on the contrary, the N-terminally tagged protein demonstrated an altered subcellular distribution. This observation can be potentially explained by the essential role that the APE1 N-terminal domain has in regulating binding activities, as extensively demonstrated by previous work from our group (7, 10, 15, 21–25) and as recently reviewed by Bañuelos and colleagues (26). Considering that APE1 altered cellular localization can be a clue to altered protein function, we proceeded with the characterization of only C-terminus-tagged clones. To assess whether the addition of the tag could alter known APE1 activities, we performed cell cycle analysis and tested cellular responses to genotoxic stimuli directly associated with APE1 DNA repair activity. All the results obtained (Fig 1E and F) led us to conclude that neither the tags nor the reduced protein expression level observed in the selected clone could significantly alter cellular homeostasis and DNA damage response. Following the characterization of this newly developed cellular model, we wanted to test the feasibility of tracking, in real time, the dynamic movements of APE1. To do so, we treated living cells with MMS and CDDP (Fig.2A and B). As expected, we observed APE1 nucleolar depletion upon CDDP treatment and, interestingly, no alteration in APE1 nucleolar localization was observed upon MMS treatment, confirming our previous observations in an overexpressing HeLa cancer cell line (24). Considering the established role of different platinum-based chemotherapeutic agents, including CDDP, in inhibiting RNA Pol I transcriptional activity (64–67), we wanted to assess if rRNA neo-synthesis was required for APE1 nucleolar localization. To test this hypothesis, we treated the cells with CX5461 and observed APE1 re-localization (Fig. 2C). From these results, we deduce that APE1 nucleolar localization depends on active rRNA transcription, a finding in agreement with our previous observations in overexpressing cancer cell lines (24).

Recently, Yan and colleagues showed that APE1 was able to undergo phase separation promoting ATR-Chk1 nucleolar condensate formation, emphasizing that the APE1 IDR was essential for condensate formation (40). To the best of our knowledge, the investigation is subject to two limitations: i) cellular experiments were performed under overexpression conditions using cancer cell lines transfected with ectopic APE1-expressing plasmids and ii) *in vitro* droplets were conducted using extremely high, non-physiological, concentrations of APE1 (550 μM). To overcome these issues, we generated the mESC cell model described above showing that APE1 is accumulated in rRNA transcription-dependent nucleolar condensates under physiological conditions. Considering that in human cells, we can find from 0.35 to 7×10^6^ molecules of APE1 (9) and that human cells have a volume spanning from 1,198 μm^3^ up to 2,425 μm^3^ (86), we estimated that APE1 protein concentration in human cells can range from about 0.2 to 10 μM, suggesting that the protein concentration used by Yan and colleagues might not represent an ideal *in vitro* conditions, possibly resulting in artifacts due to the elevated protein concentration. Unfortunately, the protein concentrations used in this work could severely affect the protein’s ability to undergo phase separation (35, 87, 88), representing an *in vitro* condition poorly relevant with respect to the cellular context for the reasons described above.

Besides observing that APE1^WT^ can form droplets at a concentration that can be considered close to physiological, we also observed that mutations to or deletion of the APE1 N-terminus IDR can change APE1 droplet formation. Since the role of RNA in influencing phase separation and droplet-forming abilities of several proteins is widely established in the scientific community (44), we wanted to assess its effect on APE1 condensate forming ability. Yan and colleagues stated that at the high concentrations used to perform the experiments, APE1 droplet forming ability was not dependent on DNA or RNA, supported by experiments that treated the samples with DNAse I and RNAse A. These indirect observations could be improved in order to evaluate the true contribution played by sequence- and structure-specific oligonucleotides that can remain in the samples upon enzymatic-(DNAse and RNAse) treatments. For this reason, we added RNA extracted from cells to our droplet solution and observed that RNA strongly affects the droplet-forming ability of not only APE1^WT^ but also, to different extents, of all the other APE1 mutants tested. Considering that: i) total RNA extracts are usually enriched (more than 90%) in rRNA; ii) both rDNA genes and rRNA have been demonstrated to be highly enriched in Gs and to form G4 structures (49, 50); iii) G4s seem involved in several biological processes including a newly discovered association with condensates (53, 54, 89); and iv) APE1 is able to bind to G4 structures not only in DNA (27, 29) but also in RNA (90), we evaluated the influence that rG4s may have on APE1 ability to form *in vitro* droplets. Interestingly, our results clearly indicate that APE1’s ability to form *in vitro* droplets and, therefore, its potential to form condensates within the cellular environment, is strongly promoted by rG4, but not by unstructured rPoly-U sequences, even when compared to APE1 natural substrates containing specific lesions (DNA containing AP sites).

Overall, the results of this study suggest that the ability of APE1 to form *in vitro* and *in vivo* condensates is a physiological mechanism to compartmentalize the protein within cells that strongly depends on specific RNA structures. To this end, the ability of APE1 to recognize and process RNAs containing both AP and 8-oxo-guanine sites, alone or in combination with additional proteins through condensation mechanisms, may represent an interesting future subject of investigation. These future studies, aiming at defining the regulatory functions of rG4s present in nuclear RNAs and understanding molecular mechanisms in RNA quality control, will open new opportunities for developing novel specific anticancer strategies.

## Materials and Methods

### Cell lines and culture conditions

V6.5 murine embryonic stem cells (mESC) were taken from a male individual derived from a and were kindly provided from the Jaenisch laboratory of the Whitehead Institute. mESC were cultured at 37°C, 5% CO_2_ in a humidified incubator on tissue-culture plates covered by 0.2% gelatin (Sigma, G1890) in 2i medium with LIF, made by 960 ml DMEM/F12 (Life Technologies, 11320082), 5 ml N2 supplement (Life Technologies, 17502048; stock 100x), 10 ml B27 supplement (Life Technologies, 17504044; stock 50x), 5 ml additional L-glutamine (Gibco, 25030–081; stock 200 mM), 10 ml MEM nonessential amino acids (Gibco, 11140076; stock 100x), 10 ml penicillin-streptomycin (Life Technologies, 15140163; stock 10^4^ U/ml), 333μL BSA fraction V (Gibco, 15260037; stock 7.50%), 7μL β-mercaptoethanol (Sigma, M6250; stock 14.3 M), 100μL LIF (Chemico, ESG1107; stock 10^7^ U/ml), 100μL PD0325901 (Stemgent, 04–0006-10; stock 10 mM), and 300μL CHIR99021 (Stemgent, 04–0004-10; stock 10 mM).

When cells were passaged TrypLE inhibition occurred using stem cell media (SCM) made by: 500 ml DMEM KO (Gibco, 10829–018), MEM nonessential amino acids (Gibco, 11140076; stock 100x), penicillin-streptomycin (Life Technologies, 15140163; stock 10^4^ U/ml), 5 ml L-glutamine (Gibco, 25030–081; stock 100x), 4μL β-mercaptoethanol (Sigma, M6250; stock 14.3 M), 50μL LIF (Chemico, ESG1107; stock 10^7^ U/ml), and 75 ml of fetal bovine serum (Sigma, F4135). Cells were then pelleted by centrifugation at 1,000 rpm for 3 minutes and resuspended in 2i medium containing LIF. Freezing cells was performed on cells in SCM, supplemented with a 1:1 ratio of 2x freezing media: 60 ml DMEM (Gibco, 11965-052), 20 ml DMSO (Sigma, D2650), and 20 ml fetal bovine serum (Sigma, F4135).

### Generation of plasmids for endogenous tagging of APE1 in mESC

Repair templates for tagging of APE1 at endogenous loci were generated on a puC19 backbone with left homology arms (LHA) and right homology arms (RHA); primers are listed in Supp. Table 1. PCRs were assembled using Phusion Flash High-Fidelity PCR Master Mix (Life Technologies, F548S) according to manufacturer instructions. PCR products were run in agarose gel, gel purified, and assembled using Gibson Assembly Cloning Kit (NEB, E5510S) according to manufacturer instructions. gRNA-Cas9 plasmids were generated upon cleavage with BbsI restriction enzyme (NEB, R3539M) of the px330 plasmid following NEB instruction; the cleaved plasmid was run on gel and purified. gRNAs (listed in Supp. Table 1) were ordered, together with their complementary, and annealed. Annealed products were ligated to the cleaved px330 plasmid using T4 DNA ligase (NEB, M0202T) and specific buffer (NEB, B0202S) following manufacturer instructions. All primers and gRNA were checked for single occupancy in the genome using the BLAT tool available on the UCSC genome browser. All primers were purchased from Eton; plasmid sequences were either checked by Sanger sequencing (Eton), for gRNA-Cas9 containing plasmids, or by Nanopore sequencing (Plasmidsaurus), for repair templates. For practical purposes, we decided to tag APE1 with mEGFP, HA, and FLAG tags, where both FLAG and HA tags allow for protein purification or immunocapturing.

### Tagging of APE1-mEGFP endogenous cell lines

The following procedure was applied for both N- and C-terminus tagging strategies. 1×10^6^ cells were plated with SCM, serum-deprived, in one well of a 6-well plate covered in 0.2% gelatin and reverse transfected with 1.6 μg of the plasmid containing the repair template, 0.4 μg of each plasmid containing gRNA1 and gRNA2 using Lipofectamine 3,000 (Life Technologies, L3000008) following manufacturer instruction. Culturing media was changed the day after transfection with the 2i medium with LIF. After two days, cells were sorted based on mCherry^+^ (tag encoded with Cas9) to enrich the population for transfected cells. mCherry^+^ cells were expanded for 7 days before the second sorting based on the mEGFP signal. All cells mEGFP^+^ were expanded and, at a later stage, dilution plating on a 10 cm dish was performed to allow single colony picking. Single clones were picked and expanded into a 96-well plate; upon confluency, 3 replicates of the 96-well plate were made to: i) freeze down cells; ii) access the zygosity condition of the clones (primer used are listed in Supp. Table 2). Selected clones were expanded and fully characterized as described in the result section.

### Western Blot

Cells were harvested and resuspended with lysis buffer (50 mM Tris HCl pH 7.5, 150 mM NaCl, 1 mM EDTA pH 8.0, 1% Triton X-100) supplemented with 1 mM protease inhibitor cocktail and 0.5 mM phenylmethylsulfonyl fluoride (PMSF). After 20 min of incubation on ice, lysates were centrifuged at 15,000 rpm to remove debris, and whole cell extract was quantified using the BCA Protein Assay Kit (Life Technologies, 23250) following manufacturer instructions. Cell extracts were kept at –80 °C.

Equal amounts of protein were separated on 10% Bis-Tris gels (BioRad Laboratories, 3450112) and transferred to a 0.45-µm PVDF membrane (Sigma, IPVH00010). Upon transfer, membranes were blocked in 5% nonfat milk (LabScientific, M0842) in TBST (2% Tris-HCl pH 8.0, 1.3% 5 M NaCl, 0.05% Tween 20) for 30 minutes at room temperature with shaking. Membranes were then incubated with primary antibody anti-APE1 (Novus Biologicals, NB100-101) and anti-βActin (Sigma, A5441) in 5% nonfat milk in TBST. Secondary antibodies (goat anti-rabbit IRDye-800CW, Li-Cor, 926-32211, and goat anti-mouse, IRDye-680RD, Li-Cor, 926-68070) in 5% nonfat milk in TBST were used to detect protein signal, assessed using Odyssey CLX LiCOR and quantified with Image Studio.

### Cell Cycle Analysis

2×10^6^ cells harvested and resuspended in 10 ml of −20 °C cold 90%-Ethanol in PBS. Cells were then kept at −20 °C for at least 1 h. When ready, cells were centrifugated to remove Ethanol and rehydrated with 10 ml of PBS. Upon rehydration, cells were collected by centrifugation and resuspended in 2 ml of PBS containing 10 μg/ml of RNaseA (Invitrogen, 8003088) and 20 μg/ml PI (Life Technologies, P3566) for 30 minutes and then analyzed with the cell sorter. A minimum of 11,000 cells were measured per each experimental condition. Analyses of the results were performed using FlowJo.

### MTS viability assay

2×10^4^ cells were plated in a 96-well plate covered in 0.2% gelatin with 2i medium with LIF. The following day cells were treated with different concentrations of CDDP (FisherScientific, 50148565) for 24 hours or MMS (Millipore, 129925) for 8 h. 1 hour before the end of the treatment, CellTiter-Glo (Promega, G9241) was added to the media, following the manufacturer instruction, and incubated for 1 hour prior recording the absorbance at 490nm with Safire 2 – Tecan. Each recorded absorbance value was standardized with the absorbance value of the wells that contained medium only. The experiment was performed in 4 independent experiments, each time with a technical triplicate.

### Immunofluorescence and foci count

1×10^5^ cells were plated in a glass bottom 24-well plate (Mattek, P24G-1.5-13-F) pre-treated with 5 μg/ml of poly-L-ornithine (Sigma, P4957) at 37 °C for at least 1 hour followed by 5 μg/ml of laminin (Corning, 354232) overnight at 37 °C. After overnight plating cells were treated with different doses of CDDP for 24 hours or with MMS for 8 hours and fixed with 4% PFA (VWR, AA47377) for 20 minutes and stored in PBS at 4 °C. Cells were permeabilized with 0.5% TritonX-100 (Sigma, X100) in PBS and blocked with 5% IgG-free BSA (VWR, 102643-516) in PBS. Anti pS139 γH2AX (Sigma, 05-636) in 5% BSA followed by Alexa Fluor 647 goat anti-mouse (Invitrogen, A21235) were used to detect DNA damage foci and Hoechst (Thermo Fischer Scientific, 3258) was used to stain nuclei.

The cells were imaged with a 100x/1.4 NA oil immersion objective lens on a spinning disk confocal microscope featuring a Zeiss AxioVert 200M inverted stand coupled to a Yokogawa CSU-22 confocal scanhead and controlled with MetaMorph 7.10. The excitation was achieved with semiconductor lasers enclosed in an Andor ILE laser launch and the emission was recorded with a Hamamatsu Orca-ER cooled CCD camera. The fluorescence from Hoechst, GFP and Alexa647 was excited with 405nm, 488nm and 642 nm laser lines and the emission from each channel was detected with 450/50nm, 525/50nm and 700/75 nm filters, respectively. The digital resolution of the images was 0.0572 um/pixel and z-stacks were acquired with an axial step size of 0.5 um.

Z-stack TIF images of γH2AX and nuclear Hoechst stain were segmented and measured using the following approach. Nuclei were segmented using the Cellpose algorithm (91) on maximal projections (z-axis) of the 405 nm Hoechst channel images. Channel images of γH2AX foci were segmented and measured using the Laplacian of Gaussian (LoG) filter. Images were maximally projected along the z-axis and subjected to the gaussian_laplace filter from the python scipy package (sigma = 3). To identify foci, an automatic threshold was then set as 1.5 standard deviations above the mean value of the LoG image. The image was binarized by this threshold and subjected to morphological opening to remove noise with a 3x3 selection element filled with 1’s. Foci were identified by the label function and measured by the regionprops function from the scikit-image python package. Only foci that localized in the nucleus (via the nuclear segmentation described above) were measured. The size and mean intensity of the foci were tabulated for analysis.

### Live cell imaging and analysis

1×10^6^ cells were plated in a glass bottom 6-well plate (Cellvis, P06-1.5H-N) pre-treated with 5 μg/ml of poly-L-ornithine (Sigma, P4957) at 37 °C for at least 1 hour followed by 5 μg/ml of laminin (Corning, 354232) overnight at 37 °C. Cells were imaged, at least 5 hours after being plated, with an Andor Revolution Spinning Disk Confocal (Oxford Instruments) microscope equipped with a Yokogawa CSU-X1 scanhead attached to a Nikon Ti-E inverted stand. Images were recorded with an Andor Zyla 5.5 sCMOS camera and a Nikon Plan Apo 100x/1.4 NA Ph3 oil immersion objective lens, delivering a digital resolution of 0.070 um/px. Phase contrast microscopy with a Phase 3 mask was used for nucleolar visualization and the tagged APE1 fluorescence was excited with a 488 nm laser (Andor ILE) and detected through a combination of a 405/488/568/647 dichroic mirror and a 525/50nm emission filter. The microscope was controlled with Metamorph 7.8 (Molecular Devices, LLC.) acquisition software.

An image was recorded before the treatment and then cells were treated either with 50 μM CDDP, 0.5 mM MMS, or 1 μM CX5461 and imaged in a time series to monitor the effects. For CDDP and MMS, the time series of the treatment effects started after 10 minutes from the treatment time and cells were imaged for a total of 6 hours with a 5 min interval. The time lapse for the cells treated with CX5461 started 6 minutes after the treatment time for a lapse of 2 hours with an interval of 3 minutes. The cells were kept at 37 °C, 95% humidity, and 5% CO2 with a stage-top incubation system (Pathology Devices). Perfect Focus was engaged throughout the time course of the experiment to maintain optimal focusing of the cells. Time lapse fluorescence images were flat-field corrected prior to any further processing.

The effect of each treatment was evaluated by analyzing the nucleoli-to-nucleoplasm intensity ratio of each nucleolus over time. This required segmenting the nuclei and nucleoli for each cell, which was performed with an analysis pipeline written in Python, that combines several Ilastik (92) projects for pixel classification and object detection.

Nuclei segmentation masks were obtained from the fluorescence channel and applied to the phase contrast images to isolate the nuclear bodies from the cytoplasm and other structures present in the images. Then a pixel classifier was trained to detect the nucleoli from the masked phase contrast images and an object classifier was trained to exclude dividing and dying cells from the analysis. The segmentation masks of the nucleoli were used to calculate their mean fluorescence intensity over time and the resulting masks from the subtraction of the nucleoli and the nuclei masks allowed to quantify the intensity in the nucleoplasm. The quantification was performed on the fluorescence images, where each nucleolus was associated to its corresponding nucleoplasm parent body to calculate the nucleoli to nucleoplasm intensity ratio.

The statistical significance of the change in the nucleoli-to-nucleoplasm fluorescence intensity ratio between the control and the treated cells was evaluated using a 2-way repeated measurements ANOVA, being time and group the independent variables. Tukey HSD was used for postdoc analysis to identify the timepoint where the effect of the studied drugs became significant.

### APE1 protein purification

DNA encoding the genes of interest (APE1^WT^, APE1^NΔ33^, APE1^K4pleA,^ and APE1^K4pleR^) were cloned into a MAT-tag backbone (23) at the C-terminus of the protein. The base vector was engineered to include a linker and mEGFP tag in between the APE1 protein and the tag. Gibson Assembly kit was used to insert these sequences in-frame. All expression constructs were sequenced through Plasmidsaurus to ensure sequence identity and proper in-frame addition of the different components. For protein expression, plasmids were transformed into LOBSTR cells (gift of Chessman Lab) and grown as follows. A fresh bacterial colony was inoculated into 200 ml LB media containing ampicillin and grown overnight at 37 °C. 30 ml of the overnight culture was diluted in 500 ml prewarmed LB with freshly added ampicillin and grown for 2 hours at 37 °C. IPTG was added to 1 mM and growth continued for 3 to 4 h. Cells were collected by centrifugation at 3,000 rpm in Sorval RC 6+ for 10 minutes and stored frozen at −80 °C. Pellets were resuspended in 15 ml of Lysis buffer (50 mM Tris HCl pH 7.5, 500 mM NaCl) containing Complete EDTA-free protease inhibitor (Roche, 11873580001), and sonicated 10 cycles of 15 seconds on, 60 sec off, 30% power of (Branson 250 sonicator with microtip). Lysates were cleared by centrifugation at 12,000g for 30 minutes and added to 1 ml of Ni-NTA agarose (Invitrogen, 60-0442), pre-washed 3 times with lysis buffer. Tubes containing the agarose lysate slurry were rotated for 2 hours at 4 °C. The slurry was centrifugated at 3,000 rpm for 10 minutes in a Thermo Scientific XTR refrigerated centrifuge. The pellet was then washed and centrifugated with 5 ml of Lysis buffer addition with protease inhibitors and increasing concentration of imidazole. 15μl of each fraction were run on a 12% gel and proteins of the correct size were dialyzed with storage buffer (50 mM Tris pH 7.5, 500 mM NaCl, 10% glycerol, and 1 mM DTT). Any precipitate after dialysis was removed by centrifugation at 3,000 rpm for 10 minutes. Protein concentration was measured with a BCA Protein Assay Kit (Life Technologies, 23250).

### Probes and their annealing

THF, DNA, DNA complementary, rPoly-U, and rG4 (Supp. Table 2) single-strand oligos were purchased from IDT. Oligomers for DNA sequences were obtained from (93). rPoly-U was selected as an RNA sequence not undergoing folding, as rG4 control, whereas the sequence for the predicted rG4 was obtained using QGRS Mapper (94). 45S sequence was used to evaluate the rG4 content of rRNA. The first rG4, with the highest score, encountered a setting as threshold 20 nucleotides in length, with a minimal G-group of 2, localized in the 18S subunit. Running QGRS Mapper on the 18S we found the same sequence, just shifted by 6 nucleotides, therefore we decided to test it for the *in vitro* droplet experiment.

Upon arrival, probes were resuspended at 100 μM in RNAse-free water. All probes, besides DNA complementary, have been labeled with Cy5 at the 5’; rPoly-U and rG4 were additionally marked with a methyl-G as the last nucleotide to maintain the oligomer more stable.

Single-stranded DNA oligos were annealed (THF-DNA complementary or DNA-DNA complementary) as follows: 100 pmol of each oligo was annealed with 150 pmol of its complementary in a final volume of 40 μl in annealing buffer (10 mM Tris-HCl pH 7.5, 10 mM MgCl_2_ and 1 mM EDTA). Annealing was performed by heating at 95° C and cooling down overnight.

RNA probes (rPoly-U and rG4) were singularly diluted 1:1 in a buffer containing KCl (final concentration of 100 mM). Samples were placed in a water bath brought to 87 °C and let cool down overnight.

### *In vitro* droplet assay and analysis

APE1 proteins underwent de-salinization and were concentrated to 20 μM using Amicon Ultra (Fisher Scientific, UFC503024) using a buffer containing 50 mM TrisHCl, 10% glycerol, and 1 mM DTT, following manufacturer instruction. 10x AP-buffer (500 mM TrisHCl, 500 mM KCl, 100 mM MgCl_2_, 10 μg/ml BSA, and 0.5% TritonX-100) was diluted to a 1x solution with RNAse free water to which APE1 proteins were added to a final concentration according to the experimental setup. Total RNA used for assessing its influence on APE1 ability to form droplets in vitro was extracted from mESC using an RNA extraction kit (Invitrogen, 12183025) following the manufacturer’s instruction.

Images of APE1-GFP droplets were analyzed using a custom script in MATLAB (v. R2018b). To define droplets while accounting for inhomogeneous background intensity, adaptive thresholding was applied. Images of the APE1-GFP protein channel (488 nm) were binarized with the ‘adaptthresh’ function with a sensitivity parameter of 0.4 and a neighborhood size of 99. Following binarization, speckles in the image foreground and background were removed using ‘bwareaopen’ with a pixel size of 5. Droplet area statistics were analyzed using ‘regionprops.’ The partition ratio of APE1-GFP (488 nm images) was calculated per droplet by dividing the mean pixel intensity in each droplet by the mean intensity of pixels in the background. The partition ratios of Cy5-labeled DNA and RNA probes (647 nm images) were calculated similarly, using the binary mask from the 488 channel to define the droplets. Condensed fraction was calculated by dividing the total area of droplets in the image divided by the area of the image.

### Statistical analysis

Statistical analyses were performed by using either the Student’s t-test or the 2-way ANOVA. p < 0.05 was considered statistically significant.

## Acknowledgments

We thank Richard A. Young (Professor of Biology at MIT and member of the Whitehead Institute) for his support and hospitality in the development of this project, Cassandra Rogers at the W.M. Keck Microscopy facility for imaging support, and Patrick Autissier at the Flow Cytometry Core for cell sorting procedures.

The work was supported by: Associazione Italiana per la Ricerca sul Cancro (AIRC) with [grant number IG19862] to G. T.., through the support of the Departmental Strategic Plan (PSD) of the University of Udine-Interdepartmental Project on Artificial Intelligence (2020–25), by additional grants from the University of Udine (‘Bando Ricerca Collaborativa’ granted by European Community -NextGenerationEU) and from the Consorzio Interuniversitario Biotecnologie -C.I.B. – (MUR-PRIN2022: “L’INNOVAZIONE DELLE BIOTECNOLOGIE NELL’ERA DELLA MEDICINA DI PRECISIONE, DEI CAMBIAMENTI CLIMATICI E DELL’ECONOMIA CIRCOLARE”) to G.T.

## Author Contributions

G.T. together with G.A. conceived and designed the study and supervised the experiments contributing to the interpretation of the results; G.D. performed the majority of the experiments, analyzed the data, and critically contributed to the interpretation of the results; N.M.H. performed the protein expression and purification for *in vitro* droplets; K.J.O., J.E.H. and A.M.V. performed the quantification analyses; J.M.P. contributed to the strategy and generation of the endogenously tagged cells; G.D., G.A. and G.T. mainly wrote the manuscript; N.M.H., K.J.O., J.M.P., J.E.H. and A.M.V provided critical comments and suggestions and contributed to the interpretation of the results. All authors critically read and approved the final version of the manuscript.

## Competing Interest Statement

The authors declare no competing interests.

## Supplementary Information

## Supp. Figures

**Supplementary Table 1.**
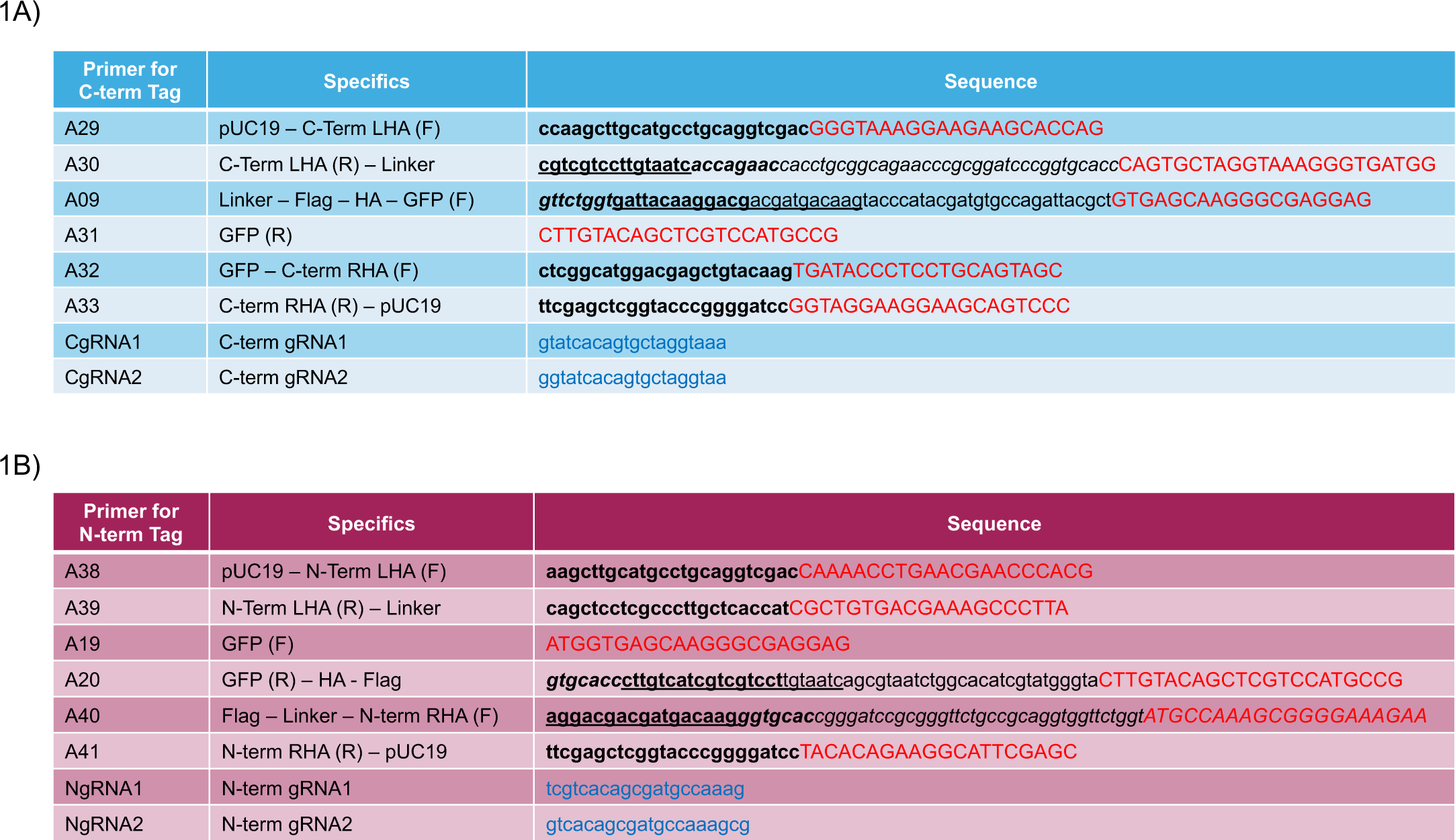
List of primers used to generate the endogenously tagged APE1 cell line at the C-terminus (A) and at the N-terminus (B). Sequences in red capital letters were used as primers for PCR; lowercase, bold, letters were used as overhangs for the Gibson assembly of the plasmids; italic lowercase letters highlight the linker sequence used to space from APE1 sequence to the tag sequence; underlined, lowercase letters mark the addition of the Flag tag whereas lowercase letters mark the addition of the HA tag; guide RNA (gRNA) sequences are reported in blue. A29-A30 and A32-A33 were used to generate PCR-amplified LHA and RHA for the C-terminus tagging location, similarly, A38-A39 and A40-A41 were used for the Nterminus homology arms. A09-A31 and A19-A20 primers were used to amplify mEGFP from a plasmid for C-terminus and N-terminus respectively.

**Supplementary Table 2.**
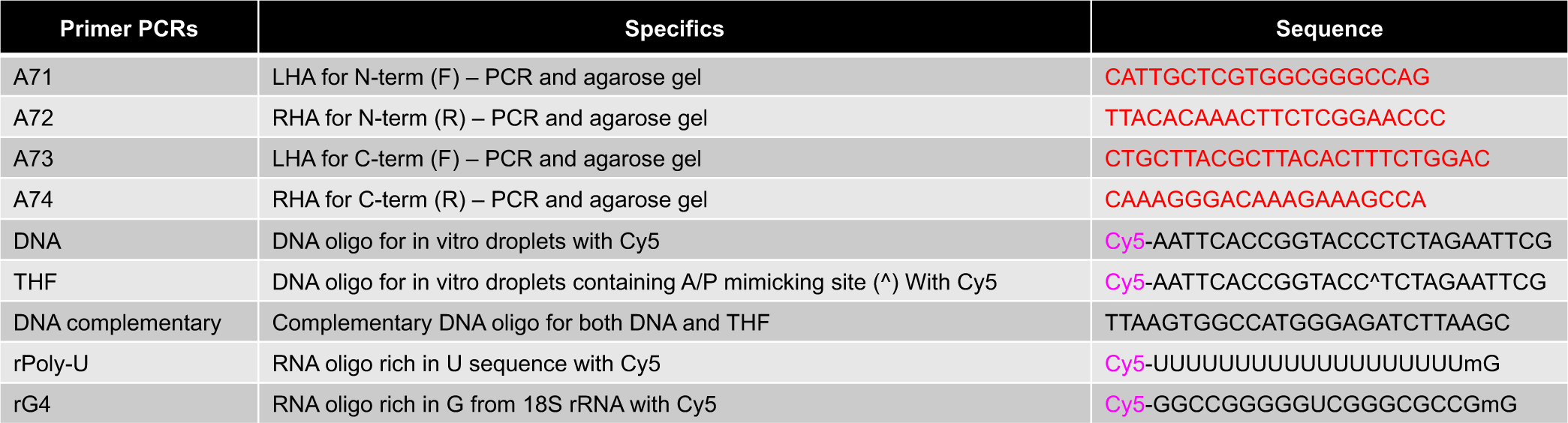
List of primers used to perform PCR from genomic DNA to validate the hetero-/homo-zygosity state of the cells upon electrophoretic agarose gel assay (A71 and A72 for the N-terminus tag, A73 and A74 for the C-terminus tag); DNA sequences used for *in vitro* droplet assay: DNA and THF, both labeled with Cy5 fluorophore at the 5’ end of the sequences, complementary sequence to generate the double-strand DNA; RNA sequences for *in vitro* droplet assay: rPoly-U and rG4, both labeled with Cy5 fluorophore at the 5’ end of the sequences with the addition of a methyl-G at the 3’ end of the sequence.

**Supplementary Fig. 1.**
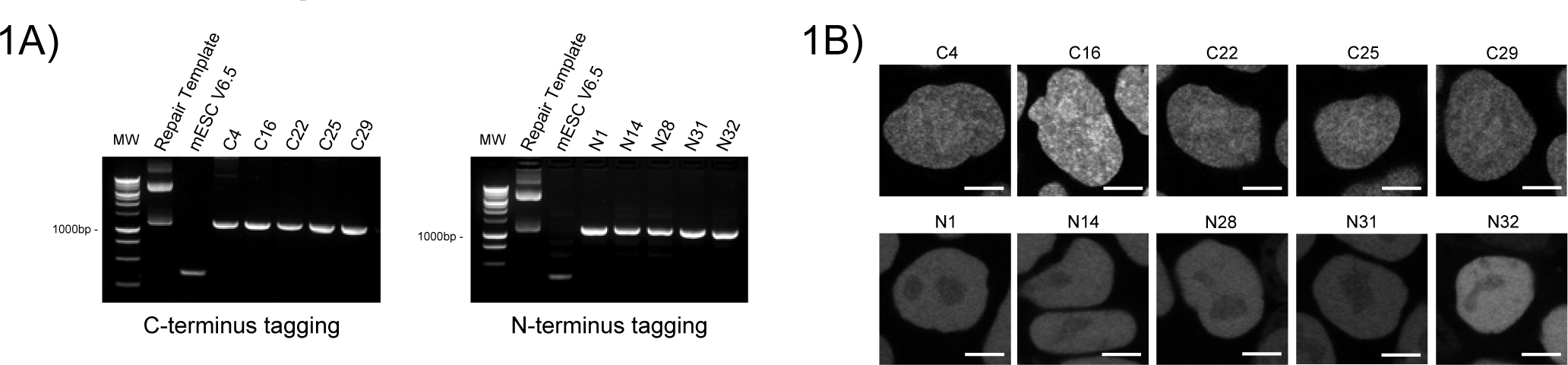
Results of the agarose gel electrophoresis performed on PCR products to validate the homozygosity of the clones (A). Live cell imaging of the 5 selected clones tagged at the C-terminus or the N-terminus; images taken with a 63x objective of a Zeiss LSM 980 with Airyscan 2 Laser Scanning Confocal microscope, scale bar 5 μm (B).

**Supplementary Fig. 2.**
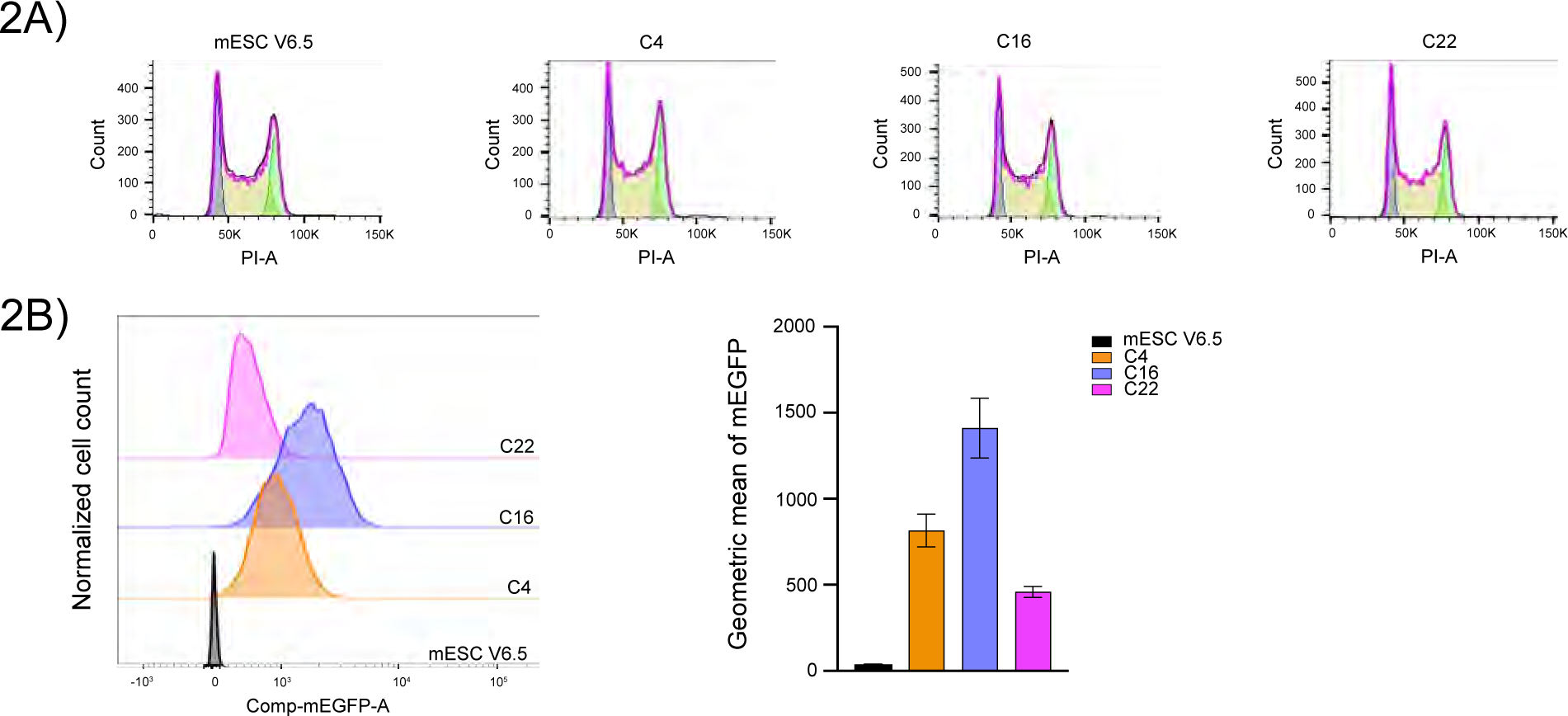
Representative result of cell sorting experiment for cell cycle analysis showing the number of cells at each cell cycle state: G0/G1, S, and G2 for wild-type cells as well as for C4, C16, and C22 clones (A). Representative result of mEGFP signal intensity and histogram showing the results of a biological duplicate (B).

**Supplementary Fig. 3.** Time-lapse video of cells treated with just media, as control, (A) or with 0.5 mM MMS (B); treated with DMF, vehicle, (C) or 50 μM CDDP (D); 0.2% acetic acid as control, (E) or RNA PolI inhibitor CX5461 (F).

**Supplementary Fig. 4.**
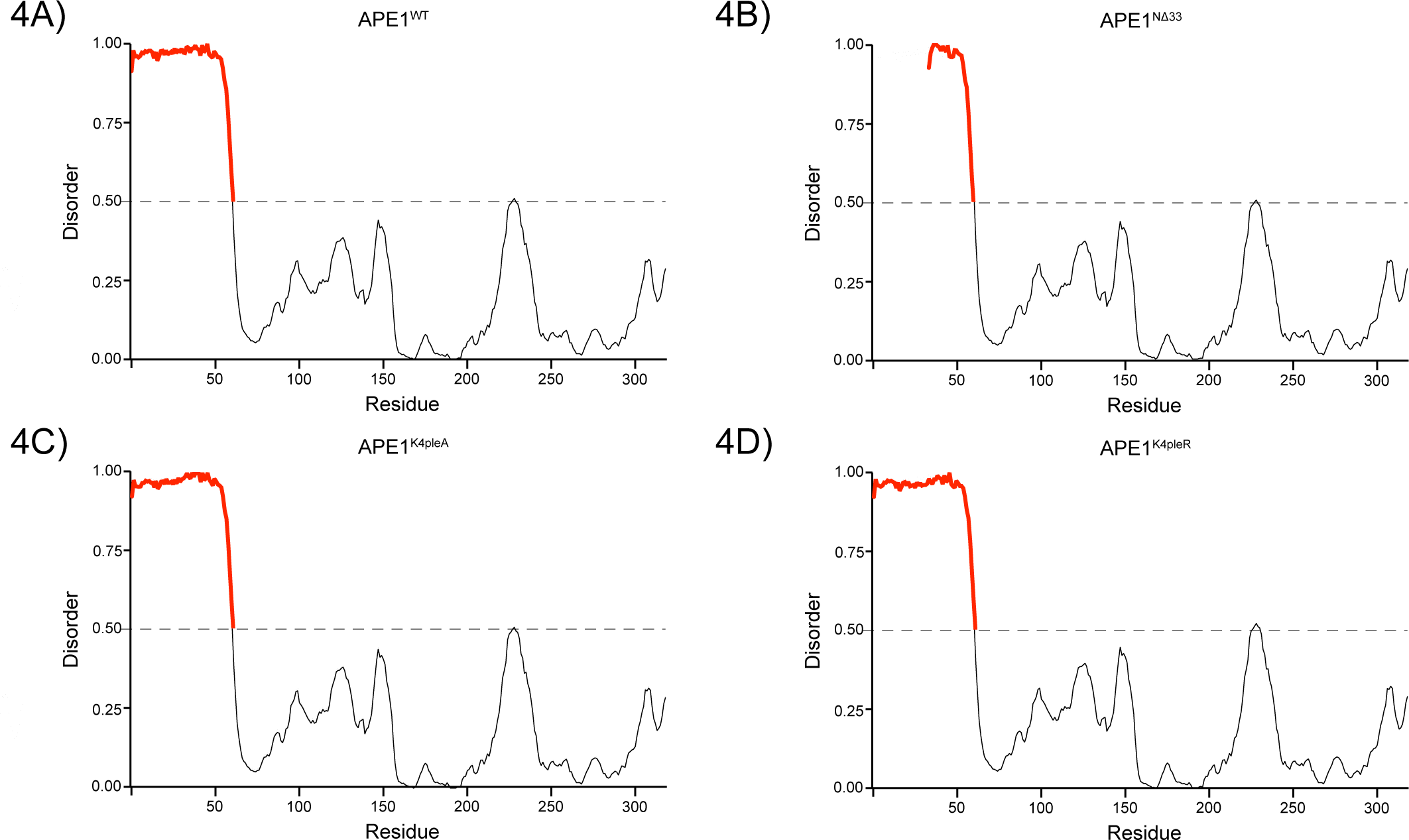
Ordered-disordered prediction of APE1 proteins using metapredict (69, 70). APE1_WT_ (A); APE1_NΔ33_ (B); APE1_K4pleA_ (C) and _K4pleR_ (D). The predicted IDR is shown in red.

**Supplementary Fig. 5.**
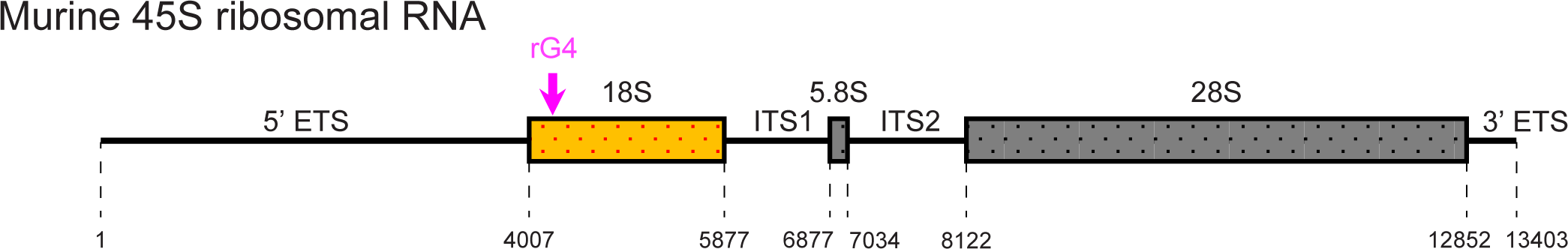
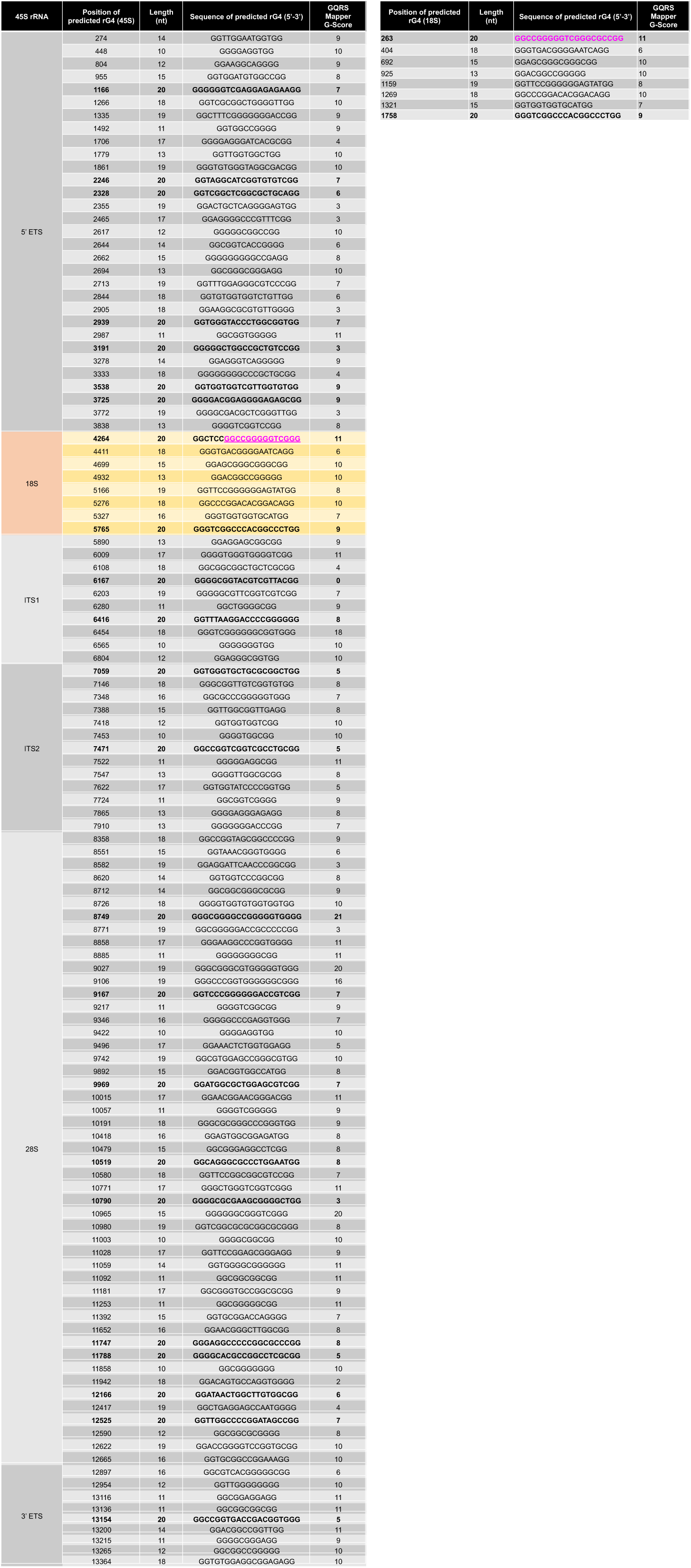
Schematic representation of the unprocessed murine 45S rRNA (top), with the annotation of its component and respective nucleotide position: 5’ ETS (External Transcribed Spacer), 18S, ITS1 (Internal Transcribed Spacer 1), 5.8S, ITS2 (Internal Transcribed Spacer 2), 28S and 3’ETS (External Transcribed Spacer); position of the selected 20-mer predicted to form G4 structure marked in magenta. Table of predicted rG4 present in the murine 45S rRNA (left) and 18S rRNA (right) with position, length, sequence, and score using GQRS-Mapper; sequences resulting from the prediction that are 20 nucleotide long are listed in bold characters; the selected sequence, predicted to form rG4 used in the *in vitro* droplet assays is highlighted in magenta.

## Notes

### Competing Interest Statement

The authors have declared no competing interest.

